# The IgCAM BT-IgSF (IgSF11) is essential for connexin43-mediated astrocyte-astrocyte and ependymal cell-cell coupling in mice

**DOI:** 10.1101/2022.06.17.496568

**Authors:** Laura Pelz, Laura Dossou, Nine Kompier, Rene Juttner, Gabrielle Siemonsmeier, Niklas Meyer, Elijah David Lowenstein, Ines Lahmann, Helmut Kettenmann, Carmen Birchmeier, Fritz Gunter Rathjen

## Abstract

The type I transmembrane protein BT-IgSF is predominantly localized in the brain and testes. It belongs to the CAR subgroup of Ig cell adhesion proteins, which have been hypothesized to regulate connexin expression or localization. Here, we studied the putative link between BT-IgSF and connexins in astrocytes, ependymal cells and neurons of the mouse. Global knockout of BT-IgSF caused an increase in the clustering of connexin43 (Gja1), but not of connexin30 (Gjb6), on astrocytes and ependymal cells. Additionally, knockout animals displayed reduced expression levels of connexin43 protein in the cortex and hippocampus. Importantly, analysis of biocytin spread in hippocampal or cortical slices from mature mice of either sex revealed a decrease in astrocytic cell-cell coupling in the absence of BT-IgSF. Blocking either protein biosynthesis or proteolysis showed that the lysosomal pathway increased connexin43 degradation. Localization of connexin43 in subcellular compartments was not impaired in the mutant. In contrast to connexin43 the localization and expression of connexin36 (Gjd2) on neurons was not affected by the absence of BT-IgSF. Overall, our data indicate that the IgCAM BT-IgSF is essential for correct gap junction-mediated astrocyte-to-astrocyte and ependymal cell-to-ependymal cell communication.

**Significance Statement:** Astrocytes regulate a variety of physiological processes in the developing and adult brain that are essential for proper brain function. Astrocytes form extensive networks in the brain and communicate via gap junctions. Disruptions of gap junction coupling are found in several diseases such as neurodegeneration or epilepsy. Here, we demonstrate that the cell adhesion protein BT-IgSF is essential for gap junction mediated coupling between astrocytes in the cortex and hippocampus.

## Introduction

BT-IgSF (<u>b</u>rain- and testis-specific <u>Ig</u> <u>s</u>uperfamily <u>p</u>rotein, also known as IgSF11) is a cell adhesion protein belonging to a small subgroup of IgCAMs consisting of CAR (<u>c</u>oxsackievirus and <u>a</u>denovirus receptor), ESAM (<u>e</u>ndothelial cell-<u>s</u>elective <u>a</u>dhesion <u>m</u>olecule) and CLMP (<u>C</u>AR-like <u>m</u>embrane <u>p</u>rotein). The group shares a similar overall domain organization with an N-terminal V-type and a C2-type Ig domain and a highly related amino acid sequence (Owczarek et al., 2023; Raschperger et al., 2004; Rathjen, 2020). Initially BT-IgSF was described as a novel IgSF member that was preferentially expressed in the brain and testis (Suzu et al., 2002). Independently of this report BT-IgSF was also found to be up-regulated in intestinal-type gastric cancers and termed IgSF11 (Katoh and Katoh, 2003). Moreover, it was also termed V-set and Immunoglobulin domain containing 3, abbreviated VSIG-3 (Harada et al., 2005; Wang et al., 2019). The genes of BT-IgSF are located of chromosome 3 and 16 in humans and mice, respectively. The cytoplasmic segment of BT-IgSF contains a PDZ-binding motif at its C-terminus that interacts with the scaffolding protein PSD95 (Jang et al., 2015). Adhesion assays with heterologous cells showed that BT-IgSF promotes homotypic cell binding (Eom et al., 2012; Harada et al., 2005).

So far, the function of BT-IgSF (IgSF11) has been studied in neurons, Sertoli and germ cells of the testes, during osteoclast differentiation and in the organization of pigment cells in fish (Ahi and Sefc, 2017; Chen et al., 2021; Eom et al., 2012; Hayano et al., 2021; Jang et al., 2015; Kim et al., 2023, 2020; Pelz et al., 2017; Singh and Nusslein-Volhard, 2015). Knockdown studies using cultured mouse hippocampal neurons indicated that BT-IgSF is implicated in synaptic transmission through a tripartite interaction with PSD95 and AMPA receptors (Jang et al., 2015). Consequently, global BT-IgSF-deficient mice displayed a moderately decreased excitatory synaptic strength in the dentate gyrus and long-term potentiation in hippocampal CA1 neurons and behavioral deficits (Jang et al., 2015; Montag et al., 2023). In another study analyzing neurons BT-IgSF (IgSF11) was found to regulate the innervation of axons of chandelier cells on initial axon segments of pyramidal neurons (Hayano et al., 2021).

In the murine testis BT-IgSF is expressed in Sertoli cells at the blood-testes barrier, a structure that opens and closes to allow the passage of germ cells. In a global knockout, the absence of BT-IgSF causes a malfunction of the blood-testes-barrier leading to male infertility due to the mislocalization of connexin43. Connexin43 was found throughout the seminiferous epithelium instead of being restricted to the blood-testes barrier as it is in wildtype animals. Therefore BT-IgSF might regulate the localization or activity of connexin43 in Sertoli cells (Pelz et al., 2017). In line with this finding is that connexin43 plays an essential role in tight junction reassembly at the blood-testes barrier during its restructuring processes (Li et al., 2010). A critical role of BT-IgSF in regulating the organization of pigment cells into stripes along the dorso-ventral or anterior-posterior body axes was observed in zebrafish and *Neolamprologus meeli* (Ahi and Sefc, 2017; Eom et al., 2012; Singh and Nusslein-Volhard, 2015). Similar irregular patterns of chromatophores were described in zebrafish with mutations in connexin 41.8 and connexin39.4, suggesting a functional link between BT-IgSF and connexins. This link might also be anticipated from the mislocalization of connexin43 in Sertoli cells of BT-IgSF knockout mice (Haffter et al., 1996; Irion et al., 2014; Pelz et al., 2017; Watanabe et al., 2006; Watanabe and Kondo, 2012). Together, these data suggest an essential function of BT-IgSF in regulating the localization or activity of connexins as it has been discussed for other members of the CAR subgroup.

To investigate the functional interaction between BT-IgSF and connexin43 further, we investigated its role in the brain on glial cells. Here, we show that BT-IgSF is strongly localized on the surface of astrocytes and ependymal cells in addition to its previously described neuronal expression (Higashine et al., 2018; Jang et al., 2015; Suzu et al., 2002). In the absence of BT-IgSF in a global mouse knockout we observed an aberrantly clustered localization of connexin43 on astrocytes and ependymal cells, and a severe reduction in connexin43 at the protein level. Consequently, dye loading experiments revealed reduced diffusion within the astrocytic network in the hippocampus as well as in the cortex of mutant mice. We discuss these findings in the context of the function of the related proteins CLMP and CAR which also affect the expression and localization of connexin43 and 45 in the smooth muscle cells of the intestine and in cardiomyocytes, respectively (Langhorst et al., 2018; Lim et al., 2008; Lisewski et al., 2008; Matthaeus et al., 2023; Rathjen, 2020).

## Material and Methods

### Mice

The global knockout of BT-IgSF (B6-Igsf11^tm1e(KOMP)Wtsi^/FGR) and its genotyping has been described elsewhere (Pelz et al., 2017). Breeding was either from tg/wt to tg/wt or tg/tg (female) to wt/tg (male). Heterozygous BT-IgSF mice did not differ from wt/wt mice (Pelz et al., 2017). Cx36-deficient mice (B6.129P2-Gjd2tm1Kwi/Cnrm; EM:00326) were obtained from the European Mouse Mutant Archive (EMMA) and genotyped as described (Güldenagel et al., 2001). Animals were housed on a 12/12 h light/dark cycle with free access to food and water. The animal procedures were performed according to the guidelines from directive 2010/63/EU of the European Parliament on the protection of animals used for scientific purposes. All experiments were approved by the local authorities (LaGeSO) (numbers T0313/97, X 9007/16, X9008/20, O 0038/08 and H0027/20). Wildtype littermates served as controls.

### Cell culture, antibodies and immunocytochemistry

Hippocampal cells were prepared from postnatal day 2 or 3 mice and cultured on poly-D-lysine coated coverslips in 24-well plates at a density of 7.5 x 10^4^ cells/ml in Neurobasal (Invitrogen) supplemented with B27 (Invitrogen) and 10% FCS (Gibco) for 10 days. Every third day half of the culture medium was removed and replaced by Neurobasal/B27 without FCS. Glial cells from postnatal day 2/3 cortices or hippocampi were prepared according to standard procedures and were maintained in DMEM supplemented with 10% FCS and penicillin/streptomycin (McCarthy and de Vellis, 1980). For immunocytochemistry cells were fixed in 4% paraformaldehyde/PBS for 10 minutes, washed with PBS/1% BSA. For stainings of cryosections (thickness 12-16 µm) by indirect immunofluorescence, adult mice were transcardially perfused with 50 ml PBS followed by 50 ml 4% paraformaldehyde. Brains were post fixed for 5 hours and then transferred into 15% followed by 30% (w/v) sucrose in PBS to obtain cryoprotection. Primary and fluorophore-conjugated secondary antibodies were applied in blocking solution (5% goat serum, 1% BSA, 0.1% Triton X-100 in PBS). Mouse monoclonal antibodies were incubated on sections using the MOM immunodetection kit from Vector Laboratories (BMK-2202). Sections or monolayer cultures were counterstained with the nuclear stain DAPI at 1 µg/ml. For antibodies see table 1.

Microscopic images were obtained at room temperature by confocal imaging using a Carl Zeiss LSM 700 Laser Scanning Microscope equipped with ZEN 2011 software and the following lenses: a Plan-Neofluar 20x/0.30 NA objective, a Plan-Achromat 40x/1.3 Oil, a Plan-Achromat 63x/1.40 NA oil objective or a Plan Apochrome 100x/1.40 oil objective (all from Carl Zeiss MicroImaging, GmbH). Figures were assembled using Illustrator CS5 (Adobe).

To quantify connexin43 clusters confocal images were taken with a 63x objective from cryosections and a 100x objective for cultured astrocytes. Quantification of connexin43 clusters was done by Fiji software setting threshold to RenyiEntropy routine; clusters were accepted between 0.05 and 1.00 µm^2^ (2-40 pixels). Primary branches of astrocytes and the density of astrocytes were counted from hippocampal slices stained by anti-GFAP and DAPI positive cells.

Analysis of localizations or co-localizations of connexin43 to intracellular compartments were done with antibodies to γ-adaptin, LAMP-1, ZO-1, GM130, to phalloidin-Alexa594 (6.6 µM; 1:500; Thermo Scientific #A12381) or to BT-IgSF (Rb96) on 4% paraformaldehyde/PBS-fixed (5 minutes on ice) on wildtype and knockout astrocyte cultures. 20 000 cells were grown on poly-D-lysine coated glass coverslips (12 mm in diameter). Assessment of cell surface localization of connexin43 cluster on knockout astrocytes was done by life staining of astrocytes with WGA-488 (1 µg/ml; ThermoFisher Scientific W11261) at 4^0^ C for 45 minutes followed by washing, PFA fixation and solubilization. Then rabbit antibodies to connexin43 were added as described above. Images were obtained by confocal microscopy using the 100x oil objective mentioned above and were analyzed using Fiji/Image J. Calculation of the Pearson correlation was determined by Coloc 2 derived intensity-based correlation analysis from regions of interest around connexin43 clusters (10 µm circles). Costes threshold regression was applied and Pearsońs R value (P) above threshold was used, data were accepted with an Costes p-value >0.95. Mann-Whitney U-test (GraphPad Prism 6) was used to compare wildtype with knockout values.

### Generation of a connexin 36 fusion protein and polyclonal antibodies to connexin36

Since several commercial anti-connexin36 antibodies were of limited use we generated polyclonal anti-mouse connexin36 antibodies in rabbits using a fusion protein that comprised the second cytoplasmic segment of mouse connexin36 (amino acid residues 99 – 197) attached to a histidine stretch. The cDNA of this segment was synthesized by Invitrogen and cloned into plasmid pMA-RQ and further subcloned into the bacterial expression vector pET-14b (Novagen/EMD Millipore). The protein was expressed in Bl21 bacteria with the addition of IPTG to a final concentration of 1 mM. Bacteria were harvested by centrifugation and frozen at 80°C. Bacterial pellets were resuspended in ice-cold lysis buffer containing 2 M Urea, 50 mM Tris, 150 mM NaCl, pH 7.4, supplemented with protease blockers (aprotinin, PMSF, leupeptin, pepstatin). Unsolubilized material was removed by centrifugation and the supernatant was precipitated by ammonium sulfate (50% saturation). The pellet was dialyzed against 10 mM Tris, pH 11 and run over an anion exchange column (DE52 Whatman). The unbound fraction was applied to an NTA column (Qiagen), washed with 20 mM imidazole and bound protein was eluted by 200mM imidazole. Purity was analyzed by 15% SDS PAGE. Rabbits were injected with 100 µg protein in Freund’s adjuvant at fortnightly intervals. The IgG fraction was obtained by protein A affinity chromatography (GE Health Care) and further purified by affinity chromatography on an affinity column containing the above mentioned connexin36 segment protein coupled to CNBr-activated Sepharose (14 mg to 2.5 g Sepharose 4B; GE Healthcare). The specificity of the affinity purified antibodies was tested on tissue extracts and cryostat sections (12 µm thick, mildly fixed with 1% paraformaldehyde/PBS for 1 minute on ice) from wildtype or connexin36-deficient mice.

### Biochemical methods

To obtain a crude membrane fraction from tissues of BT-IgSF knockout or wildtype mice of different ages (as indicated in the Figure legends), hippocampi or cortices were homogenized in 0.34 M sucrose supplemented with protease blockers [aprotinin (20 U/µl), leupeptin (5 mM), pepstatin (5mM), PMSF (1mM)]. Nuclei were pelleted at 200xg for 10 minutes and the resulting supernatant was centrifuged at 100 000xg for 10 minutes to obtain a crude membrane pellet and cytoplasmic fraction in the supernatant. Membranes were stripped with 0.1 M diethylamine (pH11.5), supplemented with protease blockers to remove peripheral membrane proteins. The membrane fraction was first solubilized in 1% Triton X100 and unsolubilized material was removed by centrifugation. The pellet was then solved in 1% SDS in PBS supplemented with protease blockers and unsolubilized material was again removed by centrifugation (Musil and Goodenough, 1991). Protein concentrations were determined using the Bradford assay (Bio-Rad #500-0006) and spectrophotometric measurements. Equal amounts of proteins were loaded on SDS-PAGE for Western blotting which was controlled by Ponceau protein stain and housekeeping proteins such as GAPDH, heavy chain of clathrin, or α-tubulin. Depending on the tissue or antibody, 10, 15 or 20 µg of protein was loaded per lane. Blots to identify BT-IgSF in cells or tissues SDS PAGE was run without reducing agents. For the calculation of the molecular mass of connexin36 in neural tissues the following molecular mass standards were used (in kDa): Conalbumin, 76; BSA, 66, actin, 43; GAPDH, 36 and carbonic anhydrase, 31.

Cycloheximide chase experiments (100 µg/ml, Sigma, C-7698, dissolved in DMSO and diluted 1:500 in the incubation medium) were done with wildtype and BT-IgSF knockout astrocytes cultures for times indicated in the Figure 3G to H. 200 000 cells were grown in 24 well clusters in DMEM/10% FCS and washed two times with DMEM without FCS before the addition of cycloheximide. Chloroquine experiments (100 µM; Sigma C6628, dissolved in PBS) to inhibit lysosomal degradation were done with wildtype and BT-IgSF knockout astrocytes cultures for 4 hours in DMEM without FCS. Treatment of wildtype astrocytes with tumor necrosis factor-α (1 ng/ml; Sigma T5944) or interferon-γ (10 ng/ml; Sigma I17001) were done in DMEM without FCS for 24 hours. In these blocking experiments cells were lysed in TBS (pH 7.4) with 1% SDS and 1 mM EDTA supplemented with protease blockers and boiled in SDS-PAGE sample buffer. Equal amounts of protein were loaded on 10% SDS PAGE, blotted and analyzed with rabbit anti-connexin43 or rabbit 96 to BT-IgSF. Equal loading was controlled by Ponceau protein stain and mAb to GAPDH. Quantification of connexin43 bands was done using Image Lab software (BioRad) and analyzed by Mann-Whitney U-test and Two-Way ANOVA (GraphPad Prism 6). Total connexin43 expression was derived from the sum of bands P0, P1 and P2 (Bands P1 and P2 correspond to phosphorylated connexin43).

### RNAscope on hippocampus sections

*In situ* fluorescent hybridization was performed using the RNAscope Multiplex Fluorescent Assay from ACDbio according to the manufacturer’s instructions (Wang et al., 2012). Briefly, PFA-fixed hippocampus or midbrain sections of 20 µm thickness were obtained from P20 or P10 wildtype mice, respectively, and stored at -80°C until use. Sections were thawed at 37°C for 10 mins and post-fixed in 4% PFA in PBS for 15 minutes before washing in PBS and continuing with the manufacturer’s instructions. Protease treatment was performed using Protease IV. RNAscope probes against BT-IgSF (C1 - 451131), GFAP (C2 - 313211), vGlut1 (C2 - 416631) and GAD65/67 (Gad1: C2 – 400951; Gad2: C2 – 439371 were used.

### Dye coupling in acute brain slices

To characterize the tracer spread within coupled astrocytic networks acute brain slices containing the hippocampus or the cortex were prepared from 8–10-week-old knockout or littermate controls of either sex as described previously (Maglione et al., 2010). In brief, mice were sacrificed by cervical dislocation, decapitated and their brains carefully removed and mounted in a chamber with ice-cold bicarbonate-buffered artificial cerebrospinal fluid (ACSF), composed of (in mM): NaCl 134; KCl 2.5; MgCl_2_ 1.3; CaCl_2_ 2; K_2_HPO_4_ 1.25; NaHCO_3_ 26; D-glucose 10; pH 7.4. The buffer solution was continuously gassed with carbogen (95% O_2_, 5% CO_2_). Coronal slices of 250 µm were prepared at 4°C using a vibratome (HM 650 V, Microm International GmbH, Walldorf, Germany), and stored in ACSF at room temperature (21–25°C) for up to 5 hours.

Before dye filling, slices were incubated for 20 minutes in 1µM sulforhodamine 101 (SR-101) at 35°C to label astrocytes. Astrocytes were identified by their SR-101 fluorescence at excitation and emission wavelengths of 555 and 585 ± 10 nm, respectively, using a 60x water-immersion objective (Olympus, Hamburg, Germany). For recording and for dye loading, a patch pipette (pulled from borosilicate glass, 1.5 mm outside diameter, 0.315 mm wall thickness) was filled with a solution containing 30 mM KCl, 1 mM MgCl_2_, 0.5 mM CaCl_2_, 100 mM potassium-gluconate, 10 mM Hepes, 5 mM EGTA, 0.5% biocytin (Sigma-Aldrich) and 3 mM Na_2_ATP, pH 7.3. Lucifer Yellow (10 µg/ml; Sigma-Aldrich) was added to the pipette solution and intracellular access of the solution was confirmed by excitation at 495 nm and visualization at an emission wavelength of 510 ± 10 nm. The pipette resistance ranged from 5 – 8 MΩ. Cells were passively dialyzed via the patch pipette for 20 min. In order to confirm cell identity and vitality, membrane currents were recorded with a series of de-and hyperpolarizing voltage steps (10 mV each, filtered at 2.9 kHz) from a holding potential of -70 mV ranging from -160 to +50 mV for 50 ms), using an EPC 10 patch-clamp amplifier and TIDA 5.25 software (HEKA Elektronik, Lambrecht, Germany), as described previously (Richter et al., 2014). Capacitive transients from the pipette were compensated online via the patch clamp amplifier (C_fast_) whereas membrane capacity and series resistance (C_slow_) were not compensated. The calculated liquid junction potential of the used intra cellular solutions was – 8.858 mV using Patchers Power Tools (Mendez & Würriehausen, Göttingen, Germany) and Igor Pro 7 software (Wavemetrics, Portland, OR, USA). The calculated reversal potentials of astrocytes were corrected for the liquid junction potential. Only cells whose series resistance was not higher than 125 % at the end of the dialysis phase compared to the beginning of the recording were taken into account for the following immunohistochemical experiments and the calculation of the membrane properties. After dye loading and patch-clamp recording, the pipette was carefully removed from the cell in order to disrupt the patch.

Slices were subsequently fixed in a solution of 4% paraformaldehyde in 0.1 M phosphate buffered saline (pH 7.4) overnight at 4°C. After fixation, slices were incubated in a solution containing 2% Triton X-100 (TX-100), 2% BSA and 5% normal donkey serum (NDS) in Tris-buffered saline (TBS) at pH 7.4 for 2 h at room temperature to permeabilize and to block nonspecific binding of the primary antibodies. Biocytin-filled networks were visualized with Cy3-conjugated streptavidin (1:200; Jackson ImmunoResearch, Hamburg, Germany). In addition, rabbit anti-BT-IgSF (Rb95) and guinea pig anti-GFAP antibodies were applied to label BT-IgSF and astrocytes, respectively. The floating slices were incubated with primary antibodies for 48 h at 4°C followed by secondary antibodies and DAPI. For additional antibodies see Table S1.

Slices were rinsed and mounted with Aqua Poly/Mount (Polysciences Inc., Washington, USA). Images were acquired by a Leica DM TCS SPE confocal microscope (HC APO 20x/0.75; Leica, Solms, Germany) with Leica software (LCS Lite or LAS AF Lite, respectively). Step size between z-planes in the confocal stacks was about 1 µm, but the number of imaged planes in each stack varied from slice to slice. Images were analyzed by Fiji/Image J software, using the cell counter plugin and z-axis projection functions.

### Statistics

Results are given as mean ± standard error of the mean (SEM). Data were tested using Mann-Whitney U-test or Two-Way ANOVA using GraphPad Prism software after excluding outlier by outlier test (ROUT, Q=1%).

## Results

### Absence of BT-IgSF disrupts the expression and localization of connexin43 in the hippocampus and on cultured astrocytes

BT-IgSF was first described as an IgSF member that was preferentially expressed in the brain and testis (Suzu et al., 2002). BT-IgSF and related proteins of the CAR family of adhesion proteins have been proposed to modulate the localization of connexins (Rathjen, 2020). We tested whether the absence of BT-IgSF affects the localization of connexin43 expression on astrocytes as it was found on Sertoli cells in the testes (Pelz et al., 2017). In the brain connexin43 is known to be expressed by astrocytes and ependymal cells, but not by neurons. Connexin30 and connexin26 have also been detected in adult astrocytes, albeit at lower levels (Dermietzel et al., 1989; Nagy et al., 2004). Hippocampal sections as well as astrocyte monolayer cultures were labeled with rabbit anti-connexin43 antibodies. Microscopic images of hippocampal sections revealed an altered localization of connexin43 in the BT-IgSF knockout tissue (Figure 1A and B). In the molecular layer of knockout hippocampal tissue, a pronounced decrease in the number of cennexin43 spots (31% of the wildtype), accompanied by a marked increase in the connexin43 spot size (average of 0.19 to 0.5 µm^2^ in wildtype and BT-IgSF-/- mice, respectively) was observed in high power magnifications (Figure 1C, E and F). Similar observations on the clustering and number of connexin43 spots were made on cultured astrocytes, suggesting that BT-IgSF might control connexin43 localization even in the absence of neurons (Figure 1D, H and I). Analysis of the distribution of the number of counts versus cluster size further highlights the difference between wildtype and knockouts (Figure 1G and J). Taken together, our data indicate that BT-IgSF is essential for the correct localization of connexin43 at the plasma membrane and its clustering in astrocytes.

**Figure 1.**
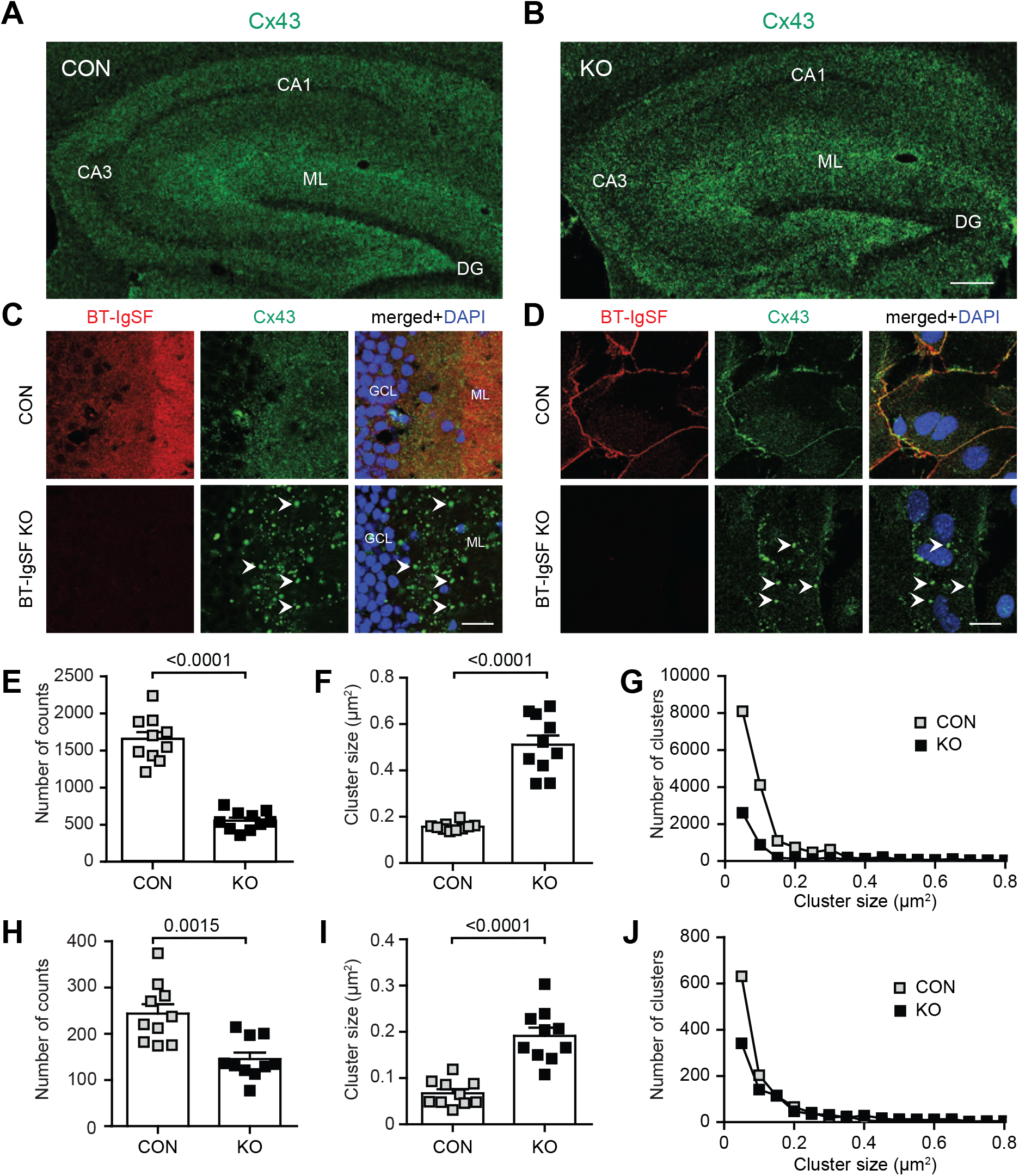
Impaired clustering of connexin43 in astrocytes in the absence of BT-IgSF. A and B) Overview of the location of connexin43 in coronal sections of the hippocampus from wildtype (A) and BT-IgSF-deficient (B) mice. 10 weeks old animals were used. CA1, cornus ammonis 1; CA3, Cornus ammonis 3; DG, dendate gyrus; GCL, granule cell layer; ML, molecular layer. Scale bar, 100 µm. C) Higher magnification of the molecular layer of the dentate gyrus from adult wildtype and knockout mice stained by mAb anti-connexin43, rabbit anti-BT-IgSF and DAPI. Arrow heads indicate large connexin43 clusters. D) Cortical astrocyte monolayer cultures from P3 wildtype or knockout mice at DIV14 stained by mAb anti-connexin43, rabbit anti-BT-IgSF and DAPI. Scale bar, 20µm. E and F) Quantification of the connexin43 spots and cluster size on astrocytes in the hippocampus. H and I) Quantification of the connexin43 spots and cluster size of connexin43 on cultured astrocytes. G and J) Cluster size distribution of connexin43 versus cluster number in the hippocampus or on astrocytes in culture (J). Scale bar in B, 100 µm; in C and D, 20µm.

We further investigated BT-IgSF expression on neural cells by immunocytochemistry of hippocampal cultures and brain sections. In monolayer cultures prepared from newborn hippocampal tissue BT-IgSF was primarily localized on the surface of GFAP-positive astrocytes (Figure 2A and B and Extended Figure 2-1 on the specificity of the anti-BT-IgSF antibody) and was not or only weakly detected on MAP2-positive dendrites (Figure 2C). However, RNAscope *in situ* hybridization experiments indicated that neurons express detectable amounts of BT-IgSF mRNA (Extended Figure 2-2). Similarly, BT-IgSF mRNA was found in GFAP–positive cells in sections of the molecular layer of the hippocampus (Figure 2E and F), which is further supported by immunohistochemistry using polyclonal antibodies to the extracellular region of BT-IgSF (Figure 2D). BT-IgSF protein expression increases as the brain matures postnatally and effective solubilization from crude membrane fractions requires ionic detergents such as SDS (Figure 2G - J).

**Figure 2.**
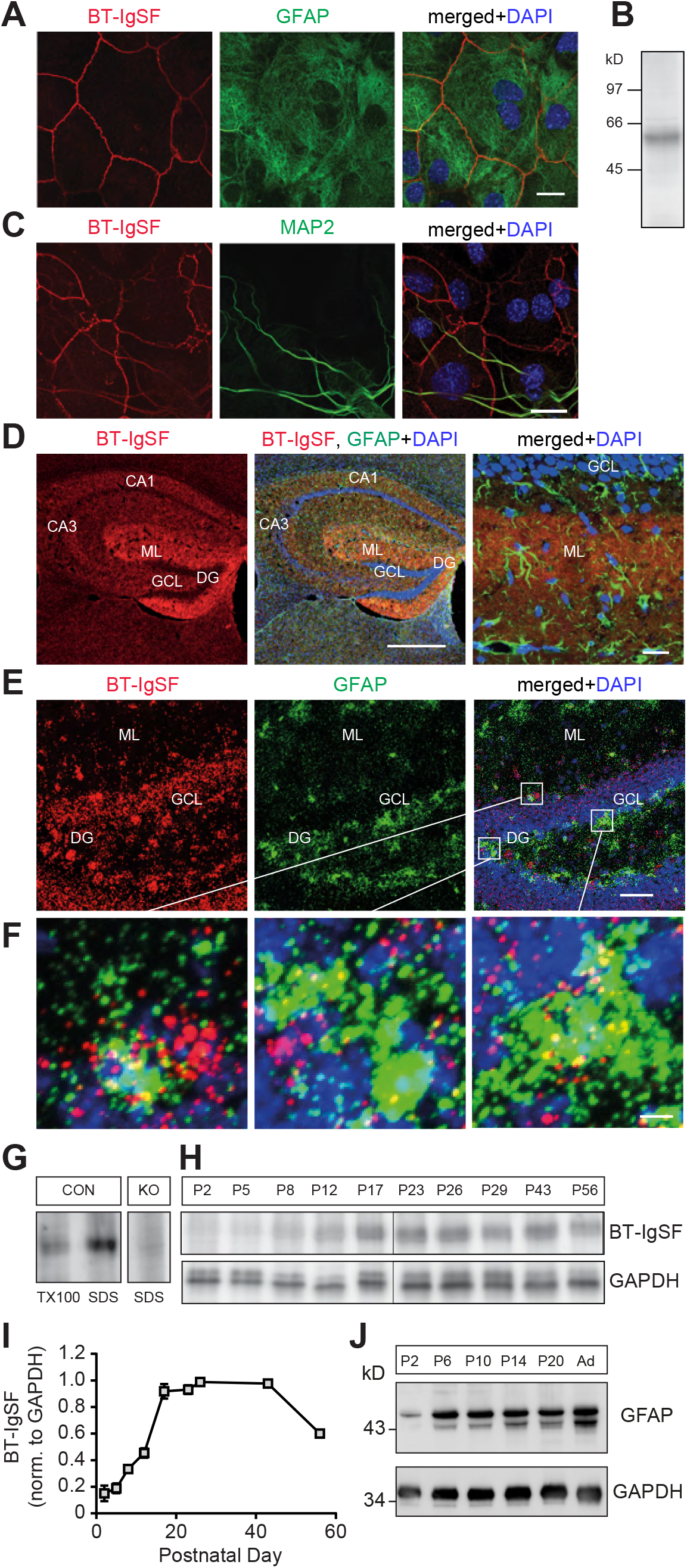
Expression of BT-IgSF on astrocytes. A) Localization of BT-IgSF in hippocampus cells from P2-old mice cultured for 10 days *in vitro* and stained by guinea pig antibodies to GFAP and rabbit anti-BT-IgSF. Scale bar, 20 µm. B) Western blot of extracts from an astrocyte culture using rabbit antibodies to BT-IgSF. Molecular mass markers are indicated at the left of the panel. C) Cultured hippocampus cells were stained by antibodies to MAP2a/b and rabbit anti-BT-IgSF. Scale bar, 30 µm. BT-IgSF is found on GFAP astrocytes but rarely detectable on MAP2a/b-positive neurons by antibodies. D) Localization of BT-IgSF in a coronal hippocampus section from an adult mouse. BT-IgSF is primarily found in the molecular layer and the subgranular zone. Higher magnification shows a widespread localization in the molecular layer probably not restricted to a specific cell type. CA1, cornus ammonis 1; CA3, Cornus ammonis 3; DG, dendate gyrus; GCL, granule cell layer; ML, molecular layer. Left and middle panel, scale bar 1 mm; right panel 20 µm. E) RNAscope of a coronal section of the hippocampus at P 20 showing expression of BT-IgSF in GFAP-positive astrocytes in the molecular layer and hilus of the dentate gyrus. Scale bar, 50 µm. F) Higher magnifications of squares as indicated in E). Scale bar, 5 µm. G) Effective extraction of BT-IgSF from tissues requires SDS. Equal amounts of crude membrane fractions from brain (P56) were extracted with 1% TX100 or SDS. For specificity of the antibody knockout brain tissue is shown. H and I) Different postnatal stages of brain extracts using SDS stained with anti-BT-IgSF are shown (n=3). BT-IgSF is primarily found at advanced postnatal stages. Anti-GAPDH indicates loading. J) For comparison GFAP expression is shown.

To analyze connexin43 in Western blots, hippocampus and cortex crude membrane fractions from wildtypes and mutants were prepared. Membranes were first treated by Triton X100 containing buffer followed by SDS of the pellet to separate soluble (primarily non-junctional) from insoluble (mainly junctional) connexin43. Significant decreases in connexin43 levels in both TX100- and SDS-containing fractions were found in the mutant (Figure 3A and B). In contrast to connexin43, connexin30, which is also implicated in the extensive network organization of astrocytes and which can also form gap junction channels together with connexin43 (Nagy et al., 2004; Willecke et al., 2002), is not reduced nor is its clustering changed in the hippocampus in the absence of BT-IgSF (Figure 3C and D). No changes in GFAP protein level were detected (Figure 3E and F) in the hippocampus of BT-IgSF mutants. This indicated that the reduction in connexin43 levels was not caused by a decrease of GFAP e.g., a decrease in the number of astrocytes (see also Figure 5K to N).

**Figure 3.**
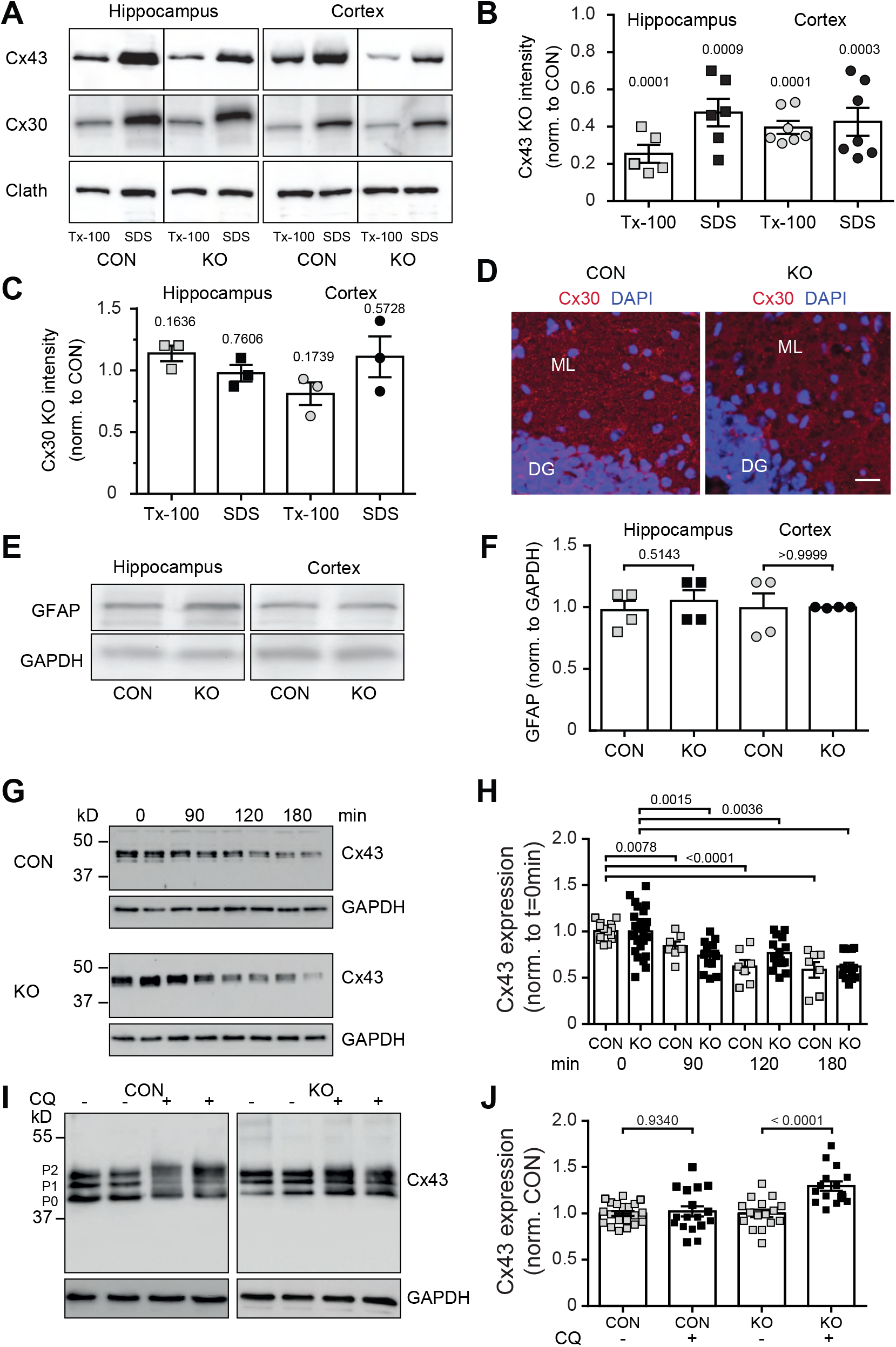
Connexin43 but not connexin30 proteins are reduced in the hippocampus and cortex, and blockers of the protein biosynthesis and proteolysis show increased degradation of connexin43 in the absence of BT-IgSF. A, B and C) Western blots demonstrating reduction of connexin43 in crude membrane fractions from 8 weeks-old hippocampi or cortices of wildtype and BT-IgSF-deficient mice. The crude membrane fractions were first solubilized in 1% Triton X100 and unsolubilized material was then solved in 1% SDS. 20 µg of protein was loaded per lane. Loading control is demonstrated by a monoclonal antibody to the heavy chain of clathrin. Quantification of band intensities are shown in B). Blot intensities of the BT-IgSF mutant were normalized to control (CON) values. p-values above the columns indicate significance to controls. For comparison quantification of Western blots of antibodies to connexin30 are shown in C. D) The localization of connexin30 in the molecular layer of hippocampi is not affected by the absence of BT-IgSF. DG, dentate gyrus; ML, molecular layer. Scale bar, 20 µm. E and F) The expression of GFAP is not reduced in the absence of BT-IgSF. Equal loading is indicated by an antibody to GAPDH. G and H) Astrocytes treated with cycloheximide (100 µg/ml) for times indicated in the Figure were lysed (1% SDS in TBS and 1 mM EDTA supplemented with protease blockers) and boiled in SDS sample buffer. Equal amounts were loaded on 10 % SDS PAGE, blotted and analyzed with rabbit anti-connexin43. Equal loading was controlled by mAb to GAPDH. Quantification of all cycloheximide experiments was done using Image Lab software (BioRad). No significance was measured between genotypes (t0 min p>0.999; t90 min p=0.1538; t120 min p=0.0938; t180 min p=0.8168; Mann-Whitney; Two-Way Anova: for time values p<0.0001; for genotypes: n.s.). I and J) Blocking of lysosomal degradation by chloroquine (CQ) (100 µM) for 4 hours in astrocyte cultures from control and BT-IgSF knockouts.

Connexin43 has a high turnover rate (Falk et al., 2016, 2014; Laird and Lampe, 2018). Therefore, the reduced expression of connexin43 might be caused by decreased biosynthesis, increased degradation or by trafficking deficits from the cytoplasm to the plasma membrane. We measured connexin43 protein levels using Western blots in a chase experiment after addition of the protein synthesis blocker cycloheximide in wildtype and BT-IgSF knockout astrocyte cultures. In both cultures, the amount of connexin43 protein significantly decreased in the presence of cycloheximide. After 180 minutes of incubation with cycloheximide the level of connexin43 protein decreased to 60 percent of baseline in both genotypes (Figure 3G and H). However, no differences between wildtype and knockout cultures were detected at different incubation periods. Linear regression analysis revealed that 50% of connexin43 would be present after 210.0 minutes and 235.1 minutes in wildtype and knockout astrocytes, respectively (p=0.8869 unpaired t-test with Welch’s correction). Interestingly, the addition of the lysosome inhibitor chloroquine for 4 hours led to an increase in connexin43 protein in knockout cultures (to 126% on average) but only sparely in control cultures (105%) (Figure 3I and J). The latter did not reach statistically significance. These data might indicate that degradation of connexin43 is increased in BT-IgSF knockout cultures. Furthermore, a change in the migration pattern of connexin43 in SDS-PAGE under chloroquine conditions was detected in astrocytes of both genotypes (Figure 3I): The connexin bands pattern became more diffuse, band P1 appeared weaker and all bands migrated at a slightly higher position which might indicate increased phosphorylation. The latter shift was stronger in the control. In summary, in the absence of BT-IgSF degradation of connexin43 via the lysosome pathway is increased which might explain the reduced expression of connexin43 in the hippocampus and cortex of BT-IgSF-/-.

To test for subcellular localization of connexin43 in the absence of BT-IgSF we compared wildtype and BT-IgSF-deficient astrocytes by staining them with antibodies specific for different compartments including LAMP-1 (lysosomal-associated membrane protein 1), phalloidin (actin cytoskeleton) GM130 (cis-Golgi marker), γ-adaptin (secretory vesicles), phalloidin (actin cytoskeleton) and ZO-1 (sub compartments such as tight junctions at the cell surface). No increased co-localization between connexin43 and any of these subcellular markers was detected in mutants indicated by a comparison of the Pearson correlation coefficients (Figure 4A to D). This might exclude that connexin43 gets retained in an intracellular compartment in BT-IgSF-/- astrocytes. Consequently, connexin43 clusters were found at the cell surface in close association with wheat germ agglutinin that was applied to living astrocyte cultures on ice (Figure 4E). Taken together these data do not support the notion that the absence of BT-IgSF reduces the expression of connexin43 in astrocytes by an impaired intracellular trafficking.

**Figure 4.**
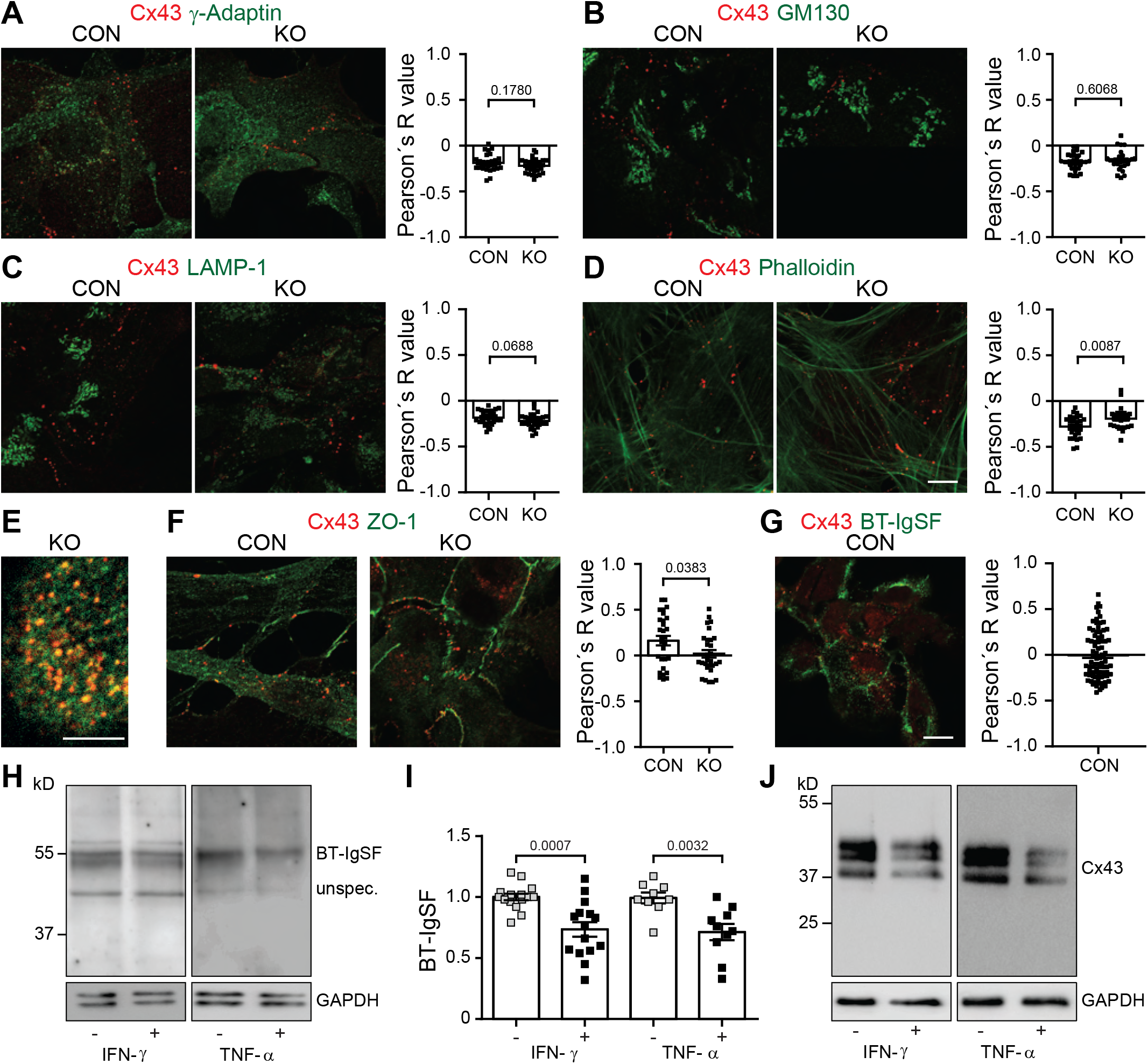
Subcellular localization of connexin43 is not impaired in the absence of BT-IgSF. A – D) The subcellular localization of connexin43 in the absence of BT-IgSF and wildtype astrocytes was analyzed by using specific antibodies to intracellular subcellular compartments. Cells were fixed with 4% paraformaldehyde/PBS (5 minutes on ice) before application of the antibodies and analyzed as described in the Materials and Method section. A) mAb to γ-adaptin, B) mAb to GM130, C) mAb to LAMP-1 D) phalloidin-594 E) Cell surface localization of connexin43 clusters on BT-IgSF-deficient astrocytes in culture. Living cells were incubated with WGA-488 for 45 minutes on ice, then fixed and permeabilized followed by staining with rabbit anti-connexin43. F) mAb to ZO-1 G) Rb96 to BT-IgSF In A to D and F) n=27-30, in G n=82 images were analyzed for the calculation of the Pearson’s R value. Scale bar in A-D and F and G, 10 µm; scale bar in E, 5 µm. H – J) Expression of BT-IgSF protein is decreased in the presence of tumor necrosis factor-α (1 ng/ml) or interferon-γ (10 ng/ml) on astrocytes in culture (H andI). Both reagents were applied for 24 hr in DMEM without FCS. For comparison reduced expression of connexin43 in the presence of tumor necrosis factor-α or interferon-γ is shown (J).

We detected minimal co-localization between connexin43 and the tight junction protein ZO-1 that was further decreased in the BT-IgSF knockout, possibly due to the fact that connexin43 was present at reduced levels (Figure 4F). In addition, BT-IgSF itself showed little co-localization with connexin43 (Figure 4G). The occasional co-localization observed might be due to the fact that all of these proteins are present at the plasma membrane, or might indicate weak and transient associations. A direct association between BT-IgSF and connexin43 could not be demonstrated since the available antibodies failed to co-precipitate BT-IgSF and connexin43.

Previously published studies reported that both the tumor necrosis factor-α and interferon-γ reduce the expression of connexin43 on astrocytes and enterocytes which in turn impairs cell-cell communication (Hinkerohe et al., 2005; Leaphart et al., 2007; Meme et al., 2006; Zhang et al., 2015, 2013). Therefore, we applied interferon-γ and tumor necrosis factor-α to wildtype astrocyte cultures. As expected connexin43 was found to be reduced (Figure 4J). Interestingly, also BT-IgSF protein level was diminished which might further strengthen our data on a functional link between BT-IgSF and connexin43 on astrocytes (Figure 4H and I).

### Reduced astrocyte-astrocyte coupling in the hippocampus and cortex in the absence of BT-IgSF

A typical feature of astrocytes in the brain is their organization in vast networks that communicate with one another via gap-junction channels formed by connexins (Giaume et al., 2010). In order to investigate the effect of the BT-IgSF ablation on astrocytic network size in the hippocampus and cortex we performed dye-coupling experiments (see schemes shown in Figure 5A to C). Astrocytes were identified in acutely isolated mouse brain slices by sulforhodamine 101 that specifically labels astroglia (Nimmerjahn et al., 2004). Individual astrocytes in the hippocampus molecular layer or cortical layers II – IV from horizontal slices were dye-loaded with biocytin via patch pipette. Diffusion of the dye throughout the astrocytic network allows for the visualization and quantification of the extent of gap-junction mediated cell-cell coupling (Figure 5D).

**Figure 5.**
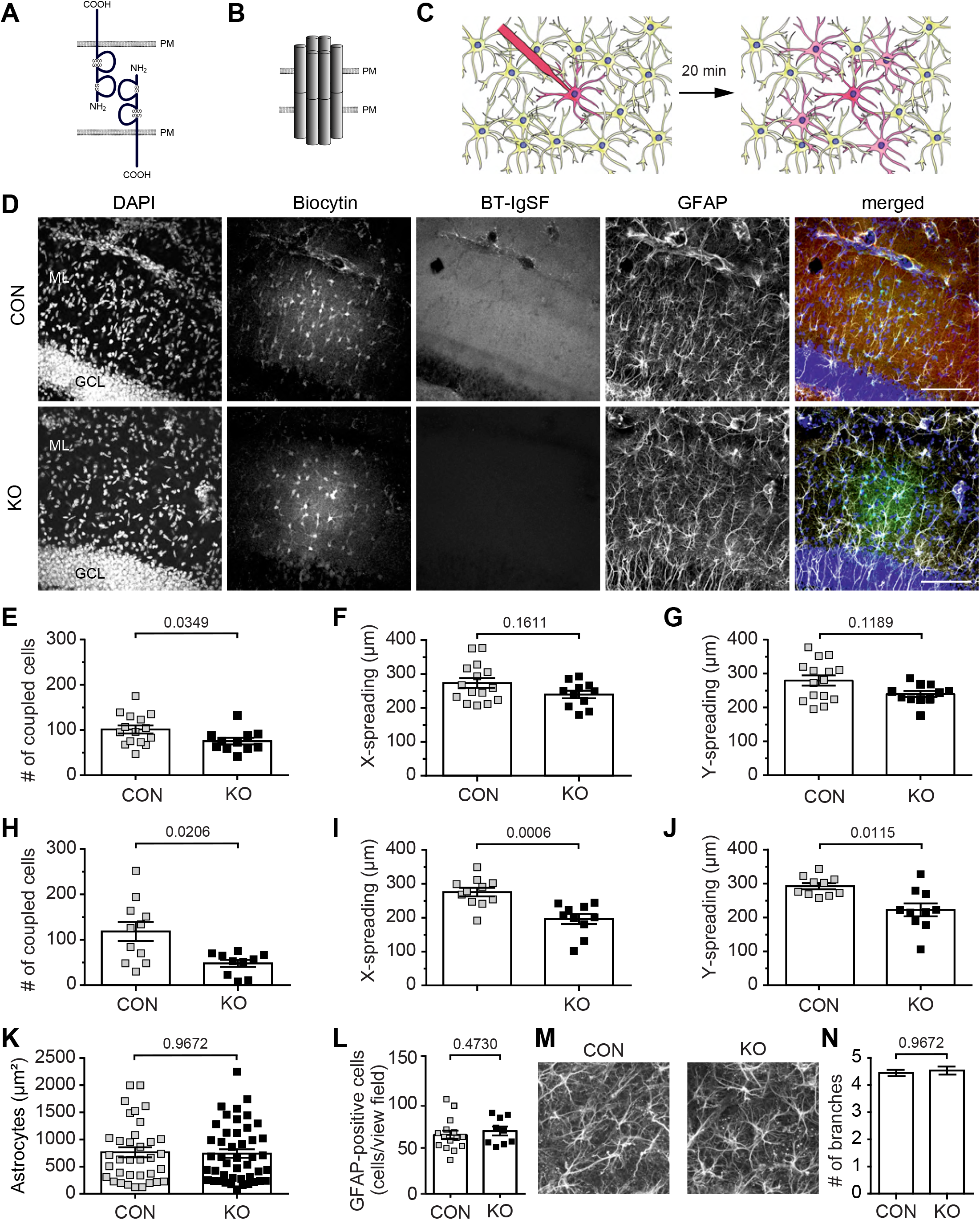
Impaired gap junction mediated coupling between astrocyte networks in the BT-IgSF knockout mice. A) Scheme of homophilic binding of BT-IgSF, B) connexin43 coupling and C) the dye filling experiment to study gap junctional communication in astrocytes. PM – plasma membrane. D) z-stack images of both genotypes of the immunohistochemical analysis for DAPI, biocytin, BT-IgSF and GFAP staining in hippocampus. The right images show an overlay of all channels. GCL, granular cell layer; ML, molecular layer. Scale bar, 100 µm. E - G) Scatter plots showing the network size (E, number of coupled cells), tracer spread alongside (F, tracer-spread x) or perpendicular to the Schaffer collaterals (G, tracer-spread y) of each dye-filled hippocampus slice. H - J) Scatter blots of dye-coupling of astrocytes in the cortex. K) Area of individual cultured astrocytes is not affected by the absence of BT-IgSF. L) Number of GFAP-positive astrocytes is not reduced in the absence of BT-IgSF. GFAP-positive cells were counted in microscopic view fields (350 µm x 350 µm) in the molecular layer of sections of hippocampi from wildtype and BT-IgSF knockout mice. 964 cells in 15 view fields and 616 cells in 9 view fields were counted for wildtype and knockout, respectively. M) Images of GFAP-positive astrocytes in the molecular layer of the hippocampus from wildtype and BT-IgSF-deficient mice indicating a similar morphology. N) Branching of GFAP-positive cells remains unchanged in the absence of BT-IgSF. Primary GFAP-positive branches were counted in the molecular layer of the hippocampus. 46 and 41 cells were counted for wildtype and knockout, respectively.

In the hippocampus as well as in the cortex, the number of coupled astrocytes per injected cell was significantly decreased in mutant animals. 16 dye-filled slices from 4 control mice and 11 slices from 3 BT-IgSF knockout mice were used for immunohistochemical analysis of hippocampal coupled astrocytic networks. On average 101±8.851 biocytin-positive hippocampal astrocytes were detected to be coupled in the wildtype, whereas in the knockout only 78±7.837 biocytin-positive astrocytes are found to form a network (Figure 4E to G). In the cortex the differences between wildtype and knockout were even more pronounced: on average 118±20.95 astrocytes in control and only 48±7.86) in the BT-IgSF knockout form a network (11 dye-filled slices from 4 control and 10 slices from 4 knockouts) (Figure 5H to J). Extended Figure 5-1 illustrates the passive membrane properties of hippocampal astrocytes in both groups over the course of the dialysis period. We observed no significant differences in the current to voltage relationship, membrane resistance or membrane capacitance (p>0.05) between genotypes or between the start and the end of the dialysis period, indicating that these factors were not responsible for the change in the extent of the astrocytic network. Similar results were obtained for passive membrane properties in cortical astrocytes. Further, neither the density of GFAP-positive astrocytes nor their complexity (arborization) was reduced in the absence of BT-IgSF, suggesting that altered morphological parameters of astrocytes are unlikely to cause the reduced cell-cell coupling in the absence of BT-IgSF (Figure 5K to N).

In summary, we detected reduced astrocyte-astrocyte coupling in the hippocampus and the cortex in the absence of BT-IgSF. We assign this to the observed changes in connexin43 expression, i.e., the reduction of the connexin43 protein levels and/or its altered subcellular localization.

### Absence of BT-IgSF disrupts the localization of connexin43 on ependymal cells

Ependymal cells are specialized glial cells that line the ventricles and form an interface between the cerebrospinal fluid (CSF) and brain parenchyma. Ependymal cells contact each other via connexin43 containing gap junctions and are implicated in barrier formation as well as the production and circulation of cerebrospinal fluid (Saunders et al., 2018). To ask whether the absence of BT-IgSF also affects the localization of connexin43 on ependymal cells, we characterized the expression of BT-IgSF in the brain by immunohistochemistry. We detected a strong localization of BT-IgSF on cells lining all brain ventricles at embryonic as well as postnatal stages, but only very weak staining in the neural tissue directly adjacent to the ependymal cell layer (Figure 6A, see also Extended Figure 6-1 for antibody specificity). Higher magnifications showed that BT-IgSF was found at the lateral and basal surfaces of ependymal cells, but not at the apical side that faces the ventricle (Figure 6B). The choroid plexus which expresses the related IgCAM CAR stained weakly for BT-IgSF (Figure 6C). Matching the deficits that we found in astrocytes, we also found increased clustering of connexin43 in ependymal cells of BT-IgSF mutants compared to wildtypes, suggesting that ependymal cell-cell coupling could also be affected (Figure 6D to G).

**Figure 6.**
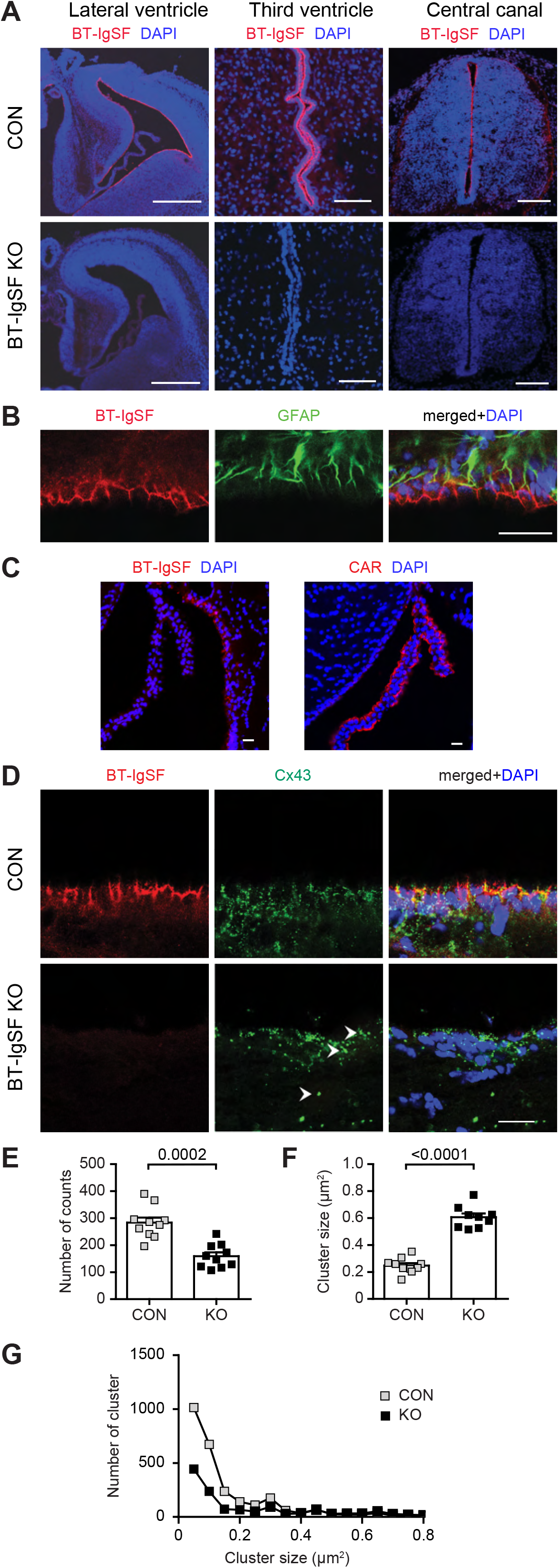
Expression of BT-IgSF on ependymal cells and impaired localization of connexin43 on ependymal cells in the absence of BT-IgSF. A) Expression of BT-IgSF in ependymal cells of the lateral, third ventricle and central canal by immunohistochemistry using an antibody to the extracellular domain of BT-IgSF. E15 lateral ventricle, scale bar 400 µm; P7 third ventricle, 50 µm; central canal of an E12.5 spinal cord, transversal section; dorsal is up, scale bar 100 µm. Red, anti-BT-IgSF; blue, DAPI. For specificity of the antibody sections of BT-IgSF knockout tissue are shown. B) Higher magnification of the ependymal cell layer showing expression of BT-IgSF at lateral and basal sides of ependymal cells in the lateral ventricle of an adult wildtype mouse. Co-localization of BT-IgSF and GFAP in the ependyma is shown. Ventricle is at the bottom of the image. Scale bar, 20 µm. C) BT-IgSF is not or only weakly found at the choroid plexus in contrast to the related CAR. Coronal sections from regions of the lateral ventricle from a 19-week-old mouse were stained with rabbit antibodies to BT-IgSF (Rb95) or to CAR (Rb80) and DAPI. D) Clustering of connexin43 on ependymal cells of the lateral ventricle from adult wildtype and BT-IgSF knockout mice. Ventricle is at the top of the images. Arrow heads indicate large connexin43 cluster. Scale bar, 20 µm. E - G) Quantification of the number of connexin43 spots, size and distribution of number versus size as described for astrocytes.

### The localization of connexin36 in neurons is not affected by the absence of BT-IgSF

In the brain, connexin43 is restricted to astrocytes, whereas connexin36 is the main connexin in neurons and is primarily expressed during embryonic and early postnatal developmental stages (Condorelli et al., 1998; Degen et al., 2004; Gulisano et al., 2000; Rubio and Nagy, 2015; Söhl et al., 1998). We therefore asked whether the absence of BT-IgSF also affects the localization of the gap junction protein connexin36 in neurons. Commercially available antibodies to connexin36 were of limited use in our hands, and we therefore generated a polyclonal antibody to the cytoplasmic stretch (residues 99 to 197) of mouse connexin36 (Figure 7A - D). Although this antibody also showed some unspecific binding in Western blots, it clearly labeled connexin36 at a molecular mass of 34 kD in wildtype but not in connexin36-deficient neural tissues (Figure 7D). The specificity of this antibody to connexin36 could also be demonstrated in sections of wildtype or connexin36-deficient brain tissue (Figure 7E). The strongest expression of connexin36 was detected in the midbrain and hindbrain at postnatal day 10, while much weaker expression levels were found in the cerebellum, basal ganglia, hippocampus and cortex. We used FISH to define the neuronal cell types expression *BT-IgSF* mRNAs, and observed in the midbrain co-expression with *vGlut1* and *GAD65*, indicating that *BT-IgSF* is expressed in both excitatory and inhibitory neurons (Figure 7F). Similar expression patterns were seen in the hippocampus (Extended Figure 2-2). Since the strongest expression of connexin36 was detected in the midbrain, we analyzed this region in more depth. No differences in the localization or expression of connexin36 were detected in the absence of BT-IgSF in the midbrain (Figure 7G and H). We conclude that BT-IgSF modulates the expression of connexin43 specifically in astrocytes and ependymal cells, but does not affect localization or clustering of connexin36 in neurons.

**Figure 7.**
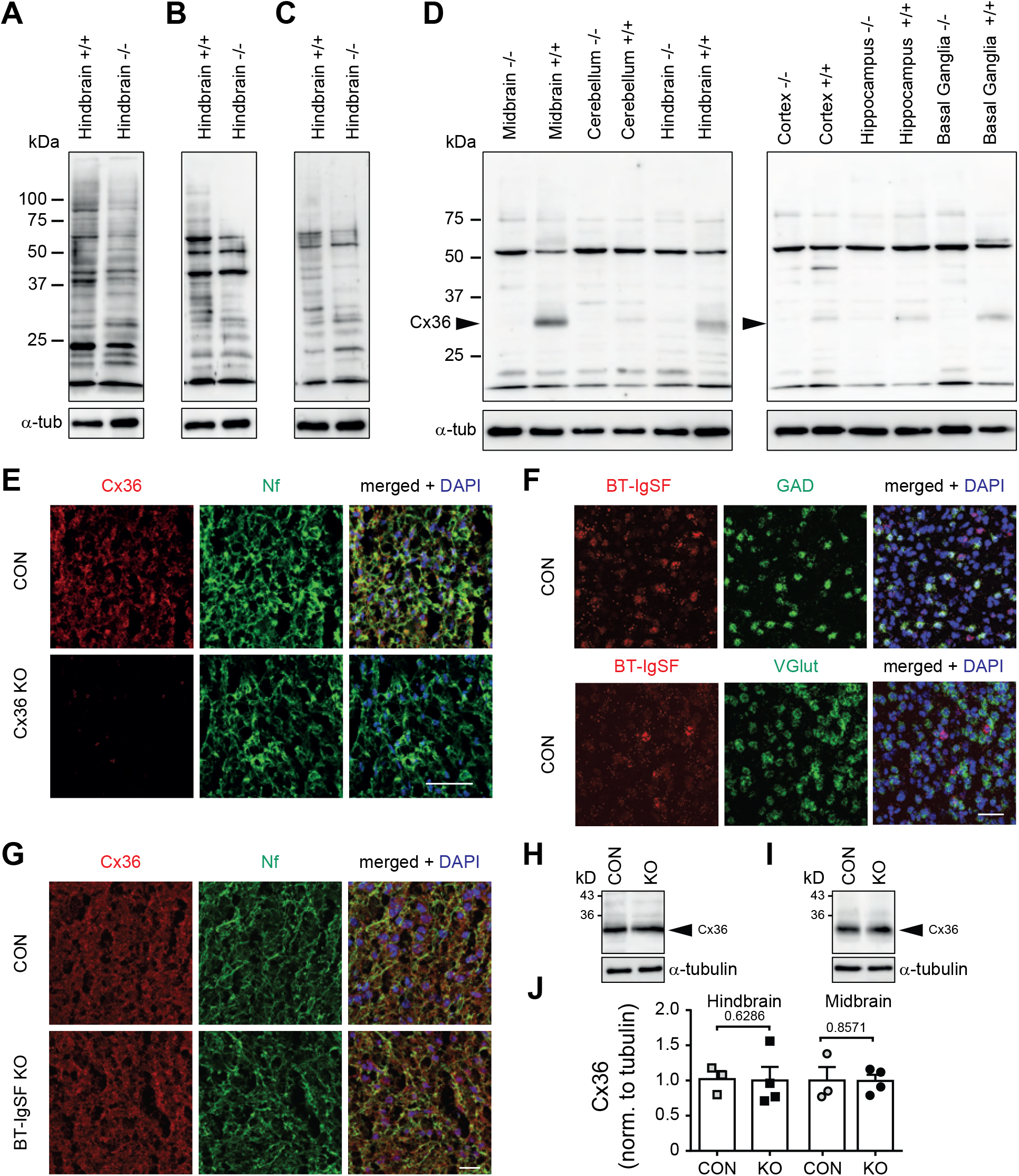
Connexin36 localization on neurons is not impaired in the absence of BT-IgSF. A - D) Specificity of antibodies to connexin36 in Western blots using crude membrane fractions from wildtype and connexin36-deficient P10 hindbrains. 12 µg of protein was loaded per lane. In A) mAb to connexin36 (sc-398063), in B) mAb to connexin36 (8F6.2), in C) rabbit to connexin36 (364600) and in D) rabbit antibody to connexin36 generated in this study (amino acid residues 99 to 197) are shown. Satisfactory specificity could only be demonstrated for antibody shown in D). A 34 kD protein (arrow) was specifically detected in wildtype but not in connexin36-deficient tissues. However, a prominent unspecific band at 52 kD was detected in both genotypes. E) Specificity of rabbit antibody to connexin36 (residues 99 to 197) in sagittal sections of P10 superior colliculus from wildtype and connexin36-deficient mice. Nf, neurofilament. F) RNAscope demonstrating neuronal expression of BT-IgSF in sagittal sections from P10 superior colliculus. G) Localization of clustering of connexin36 is not altered in P10 sagittal sections in the absence of BT-IgSF. Scale bar, 20µm. Nf, neurofilament H - J) Connexin36 protein is not altered at the protein level in the absence of BT-IgSF.

## Discussion

Here we show that the Ig cell adhesion molecule BT-IgSF is highly expressed on astrocytes and ependymal cells of the mouse brain. Our functional analysis demonstrates that BT-IgSF is essential for the correct expression and subcellular localization of connexin43 in astrocytes and ependymal cells. In the absence of BT-IgSF, the level of connexin43 protein is severely reduced and fewer but larger clusters of connexin43 are formed on the surface of astrocytes and ependymal cells in the brain. Importantly, our analysis of gap junction coupling analyzed by dye diffusion revealed a reduced astrocytic network in the cortex and hippocampus in BT-IgSF knockouts. However, this impaired coupling was not due to morphological changes of astrocytes in the absence of BT-IgSF. Astrocytes generate a complex syncytial network allowing them to interact with many neighboring cells to control a number of physiological processes like neurotransmission and disruptions of gap junction coupling are found in a broad spectrum of diseases such as neurodegeneration or epilepsy (Bedner and Steinhäuser, 2023; Huang et al., 2021; Mayorquin et al., 2018; Mazaud et al., 2021). A change in the pattern of localization or level of expression of connexin36 in neurons was not detected in the absence of BT-IgSF.

BT-IgSF is a member of a small and evolutionarily conserved subfamily of IgCAMs of which CAR was the founding member. This set of proteins mediates homotypic cell adhesion, share a common overall extracellular domain structure and a highly related amino acid sequence. A number of studies that relied on mouse mutants suggested that these proteins might undertake similar functions: they help to organize or regulate gap junctions in a variety of cell types (Falk, 2020; Rathjen, 2020). For example, in the absence of CAR, connexin43 and 45 are reduced in the heart, resulting to impaired electrical conduction at the atrioventricular node (Lim et al., 2008; Lisewski et al., 2008; Pazirandeh et al., 2011). In addition, cultured embryonic CAR-deficient cardiomyocytes showed increased calcium cycling, altered connexin43 clustering and impaired dye coupling (Matthaeus et al., 2023).Another example is CLMP: the absence of this adhesion protein impaired smooth muscle cells of the intestine and ureter. Uncoordinated calcium signaling provoked a disturbed contraction of the intestine and ureter. Notably, the level of connexin43 and 45 proteins, but not their mRNAs, was severely reduced in the smooth muscle layers (Langhorst et al., 2018; Rathjen and Jüttner, 2023). Our studies on BT-IgSF described here provide an additional example of the way this subgroup of proteins plays a part in gap junction mediated cell-cell communication. The absence of BT-IgSF also affects the expression and localization pattern of connexin43 in astrocytes and ependymal cells. This is consistent with previously published data on the mis-localization of connexin43 in Sertoli cells of the testes, which leads to infertility and the functional impairment of the blood-testes-barrier (Pelz et al., 2017). Thus, all three members of the family, BT-IgSF, CLMP and CAR, are required for appropriate gap junction mediated cell-cell communication.

### How might BT-IgSF exert its function on connexin43 in astrocytes?

Connexin43, much like other connexins, has a dynamic “life cycle” and possesses a half-life time of about 1-5 hours. Furthermore, connexin biosynthesis is complex and a plethora of proteins are known to interact with connexins during the stages of the life cycle of gap junction (Solan and Lampe, 2016). The reduced expression of connexin43 in BT-IgSF knockout astrocytes appears to be caused by increased degradation as revealed by our lysosomal pathway inhibition experiments. However, the intracellular localization of connexin43 to specific compartments was not altered, suggesting that intracellular trafficking of connexin43 might not be disturbed in the absence of BT-IgSF. The decreased amount of connexin43 might result in increased clustering at the cell surface and might therefore be a secondary effect. Interestingly, treatment of astrocyte cultures with tumor necrosis factor-α or interferon-γ caused a reduction of BT-IgSF which parallels the decrease of connexin43 by these factors. The latter finding was previously described on enterocytes of the intestine and on astrocytes (Hinkerohe et al., 2005; Leaphart et al., 2007; Meme et al., 2006; Zhang et al., 2015, 2013). Therefore, cytokines might be players that regulate the expression of BT-IgSF protein which in turn regulates connexin43 expression. However and overall, it might strengthen our proposal on a functional link between BT-IgSF and connexin43 on astrocytes. Reduced astrocyte-astrocyte coupling was also recently described in knockdown experiments of the Ig cell adhesion molecule HepaCAM (also termed GlialCAM) that, at the structural level is only distantly related to CAR subgroup members. The authors found a reduced morphological complexity, including arborization of astrocytic protrusions, and a reduction in the territory covered by astrocytes. In addition, an increased clustering of connexin43 was found in HepaCAM mutants which was accompanied by decreased dye coupling (Baldwin et al., 2021). Except for the increased clustering of connexin43 we did not detect similar morphological changes in astrocytes of BT-IgSF knockouts, nor did we find a reduction in the cell density in the molecular layer of the hippocampus. Therefore, in astrocytes HepaCAM and BT-IgSF seemingly play different roles, although in both cases their mutation resulted in impaired coupling. Furthermore, in biochemical experiments a direct association of HepaCAM with connexin43 at the cell surface was established (Baldwin et al., 2021; Wu et al., 2016). By contrast, based on our co-localization studies in cultured astrocytes, we only occasionally detected a close proximity of BT-IgSF with connexin43 at the cell surface of astrocytes, and we could not detect a direct interaction between the two proteins in co-precipitation experiments.

Co-localization of BT-IgSF with the scaffolding protein ZO-1 (Zona Occludens-1) at the cell surface was detected in Sertoli cells in confocal images (Pelz et al., 2017). BT-IgSF harbors a PDZ binding motif at its C-terminal domain and scaffolding proteins such as ZO-1 bind via a PDZ domain to most connexins (Hervé et al., 2014). Therefore, one might speculate that ZO-1 binds to BT-IgSF via its PDZ domain1, and to connexin43 via its PDZ domain 2 thus regulating connexin43 localization (Duffy et al., 2002). Furthermore, ZO-1 has been shown to control gap junction assembly and localization and influences plaque size in cell cultures (Hunter et al., 2005; Laing et al., 2005; Rhett et al., 2011), and a frameshift mutation in the connexin43 gene in human patients suffering from oculo-dento digital dysplasia disrupts the connexin43-ZO-1 interaction (Bock et al., 2013; van Steensel et al., 2005). Despite these findings in other cell types, co-localization of ZO-1 with connexin43 or co-localization of BT-IgSF with connexin43 was only rarely detected at the plasma membrane of astrocytes. Nevertheless, a transient association of these components cannot be excluded. Additional studies with mice expressing mutant versions of BT-IgSF and ZO-1 to investigate possible interactions between BT-IgSF, ZO-1 and connexin43 might clarify whether these complexes play a role for connexin43 assembly in astrocytes. Currently the question of precisely how BT-IgSF controls connexin43 localization and expression needs to be investigated further. Additional studies are needed to thoroughly understand a possible signal transduction pathway downstream of BT-IgSF that modulate the de novo incorporation or removal of connexin43 to or from the plasma membrane.

### BT-IgSF might have different functions in neurons and glial cells

Previously published knockdown studies using cultured hippocampal neurons from the CA1 region implicated BT-IgSF (IgSF11) in synaptic transmission through interactions with PSD95 and AMPA receptors in the (Jang et al., 2015). In these *in vitro* experiments, the knockdown of BT-IgSF (IgSF11) caused an increased mobility and endocytosis of AMPA receptors, suggesting that BT-IgSF is important for the stabilization of AMPA receptors in the neuronal plasma membrane. In accordance, BT-IgSF-deficient mice revealed a moderately decreased excitatory synaptic strength in the dentate gyrus and enhanced long-term potentiation in CA1 (Jang et al., 2015). Furthermore, BT-IgSF (IgSF11) was found to regulate the innervation of axons of chandelier cells on initial axon segments of pyramidal neurons (Hayano et al., 2021). To verify if our findings also apply to connexin localization in neurons, we extended our study and analyzed localization and expression of connexin36, the main neuronal connexin (Condorelli et al., 1998; Degen et al., 2004; Gulisano et al., 2000; Rubio and Nagy, 2015; Söhl et al., 1998). No differences in the localization or expression of connexin36 protein was detected in the absence of BT-IgSF in the midbrain. We conclude that BT-IgSF might have a different function in astrocytes and neurons, specifically modulating the expression of connexin43 in astrocytes and ependymal cells, but not affecting the distribution of connexin36 in neurons.

### What might be the consequences of astrocyte network disturbance for neuronal function?

Astrocytes form an elaborate network to control a number of physiological processes in the brain. Disrupted communication of astrocytes in the absence of BT-IgSF might therefore interfere with the coordination of astrocytic calcium waves or might affect synaptic activity. The close contact of astrocytic branches with synapses allows astrocytes to sense neuronal activity via their ion channels and neurotransmitter receptors (Ventura and Harris, 1999). Disruption of astrocytic networks for example by inactivation of connexin43 and 30 reduced synaptic transmission (Charvériat et al., 2017; Giaume et al., 2010; Hardy et al., 2021; Pannasch et al., 2011; Perea et al., 2009; Pereda, 2014). Accordingly, connexin30 and astrocyte-targeted connexin43 knockout mice as well as connexin43 and 30 double knockout mice display impaired performance in sensorimotor and spatial memory tasks (Lutz et al., 2009; Theis et al., 2003). Astrocyte coupling is also altered in epilepsy (Bedner and Steinhäuser, 2023; Steinhäuser and Boison, 2012). Decreased coupling among astrocytes promotes neuronal hyperexcitability and attenuates seizure-induced histopathological outcomes (Deshpande et al., 2020). Astrocytic dysfunction is implicated in a number of neurodevelopmental disorders (Molofsk et al., 2012; Tan et al., 2021). Whether the behavioral deficits observed in BT-IgSF knockouts are caused by the reduced astrocyte-astrocyte coupling or by deficits in the AMPA receptor trafficking needs further investigation using astrocyte- or neuron-specific ablation of BT-IgSF (Montag et al., 2023).

In the brain and spinal cord, ependymal cells line the ventricles and bear multiple cilia which beat in a concerted manner at their apical surface to drive cerebrospinal fluid circulation (Spassky and Meunier, 2017). This unidirectional movement might be coordinated by gap junction mediated cell-cell communication. Consistent with this hypothesis, *zebrafish* embryos injected with connexin43 morpholinos and connexin43-deficient mouse embryos exhibit a decreased number of cilia as well as diminished beating (Zhang et al., 2020).

In conclusion, in the present study we have identified BT-IgSF as a crucial molecular helper to establish correct localization of connexin43 on astrocytes and ependymal cells; in astrocytes, this is essential for effective cell-cell coupling. Whether BT-IgSF’s selective absence in astrocytes or ependymal cells is also important for neurotransmission or the coordinated movement of cilia, respectively, should be investigated in future work.

## Extended information

**Extended Figure 2-1.**
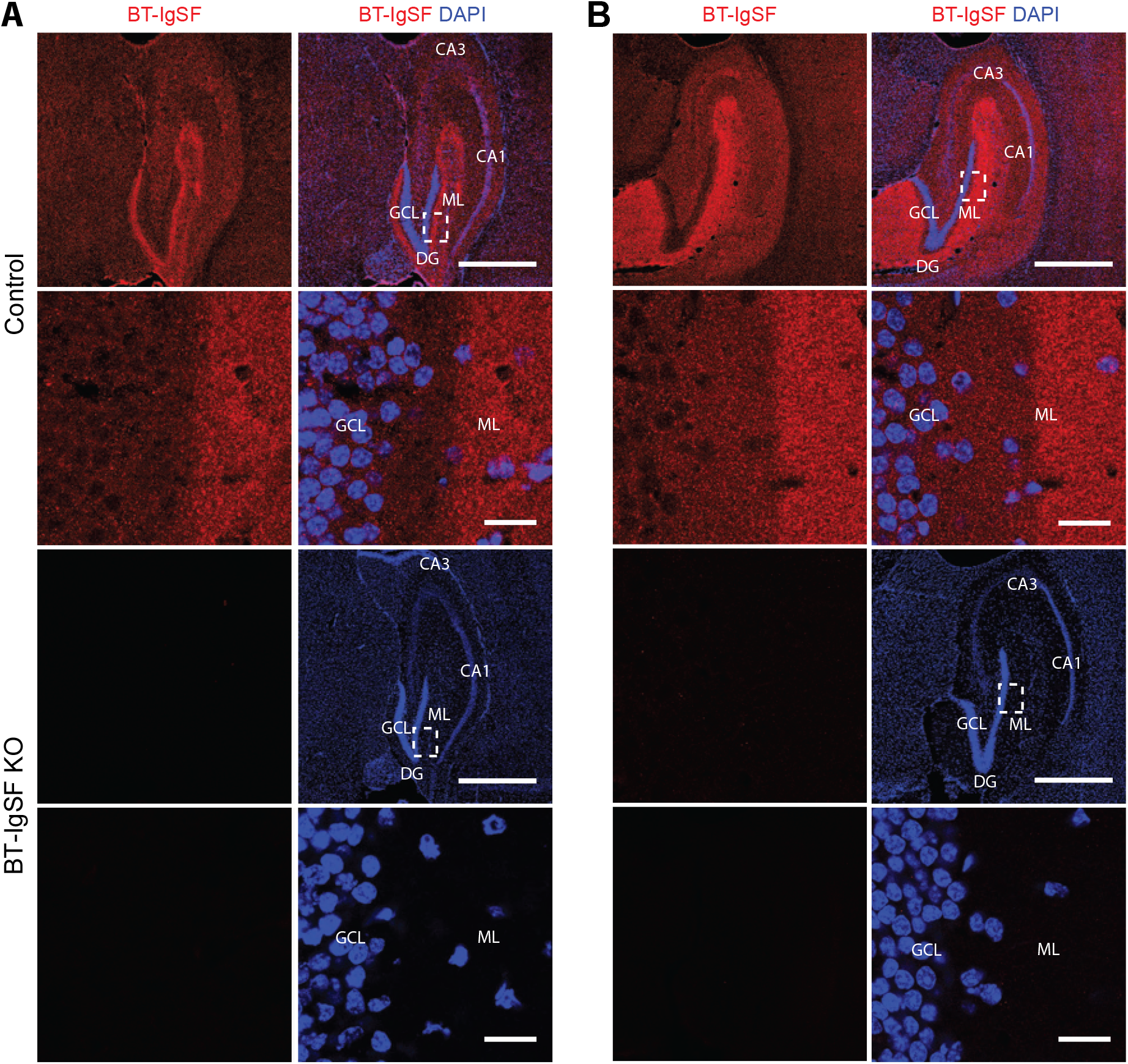
Localization of BT-IgSF in the hippocampus at P14 and P80 and demonstration of the specificity of rabbit anti-BT-IgSF. Coronal cryostat sections of the hippocampus at P14 (A) or P80 (B) were stained with anti-BT-IgSF (Rb95; 1 µg/ml). The dashed boxes indicate the position of the enlarged region shown below row 1 and 3. Absence of staining in BT-IgSF knockout tissues indicates the specificity of rabbit antibodies to BT-IgSF. CA1, cornus ammonis 1; CA3, Cornus ammonis 3; DG, dentate gyrus; GCL, granule cell layer; ML, molecular layer. Scale bar first and third row, 1 mm; scale bar second and fourth row, 20 µm.

**Extended Figure 2-2.**
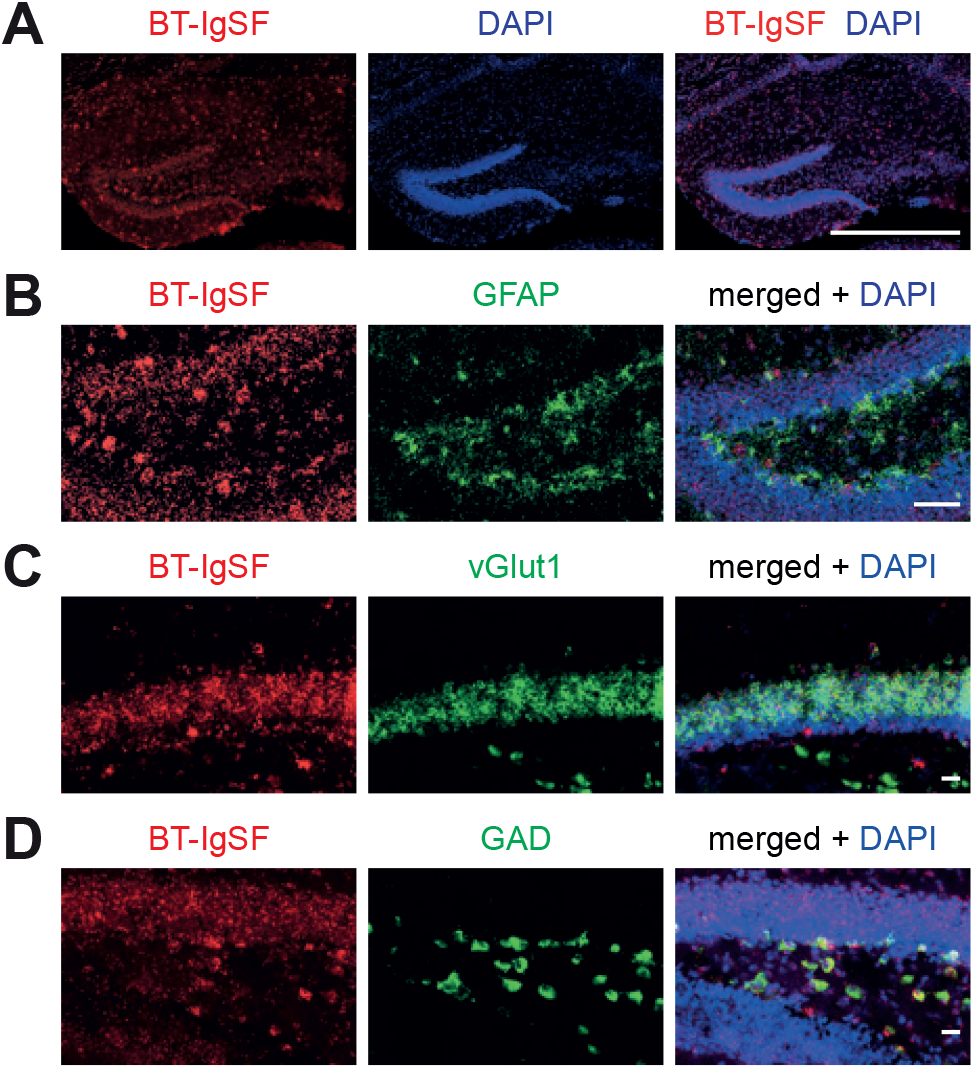
BT-IgSF encoding mRNA is expressed in excitatory and inhibitory neurons of the hippocampus. A) Overview of an RNA scope of BT-IgSF of a coronal section of a P20 hippocampus showing BT-IgSF mRNA encoding cells in all cell layers. Scale bar 500 µm. B) RNAscope of coronal sections of hippocampi at postnatal day 20 showing expression of BT-IgSF in GFAP-positive astrocytes. C) RNAscope of coronal sections of hippocampi at postnatal day 20 showing expression of BT-IgSF in vGlut1-positive neurons. D) RNAscope of coronal sections of hippocampi at postnatal day 20 showing expression of BT-IgSF in GAD65-positive neurons.

**Extended Figure 5-1.**
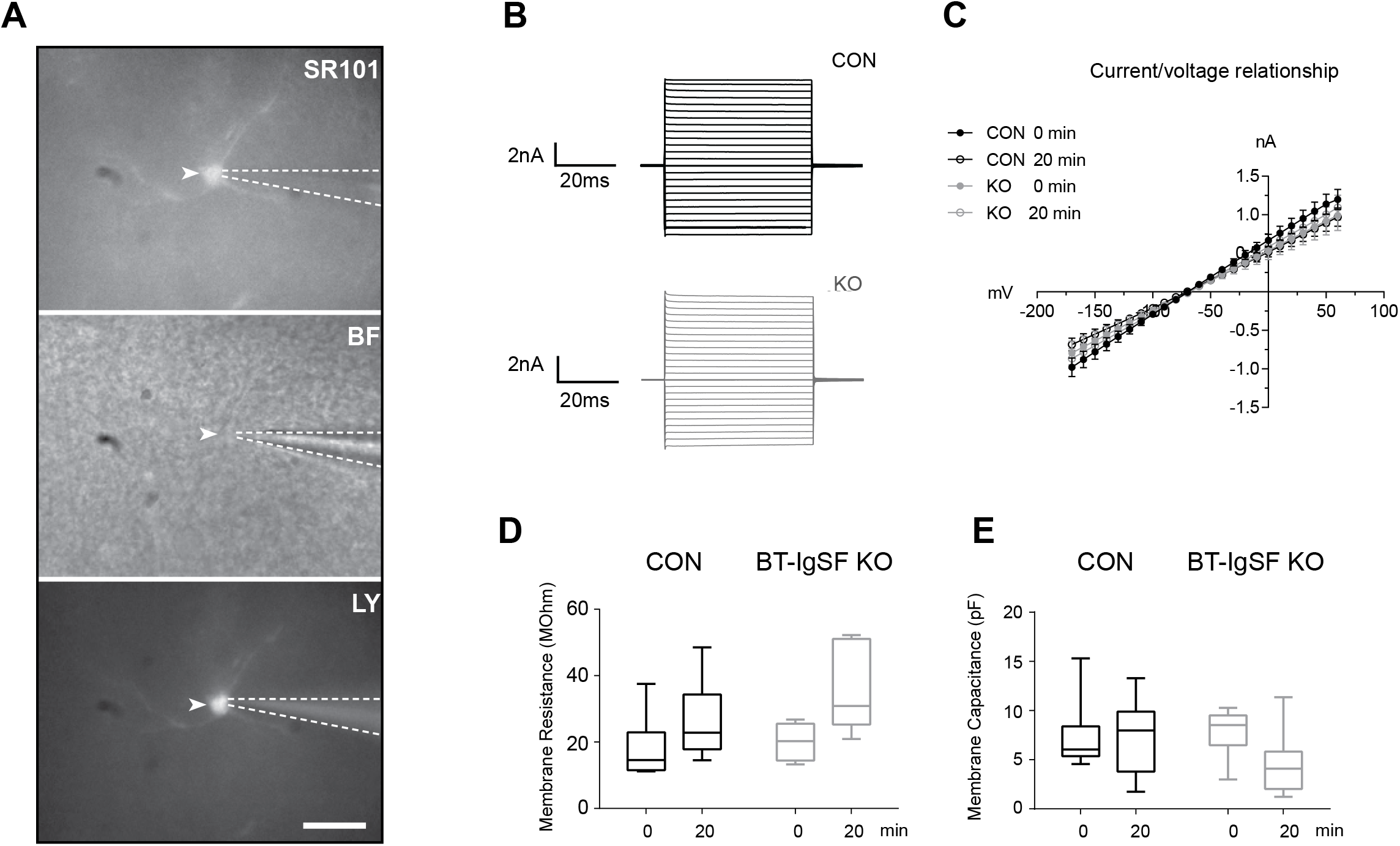
Analysis of dye spreading in astrocytes. A) Bright field and fluorescence images of a patch-clamped astrocyte in the hippocampus. *Top* Sulforhodamine (SR101) staining of the patched astrocyte. The patch-clamp pipettes outline is marked as a dotted line. *Middle* Bright field (BF) image of the field of interest. The arrow marks the position of the patched astrocyte. *Bottom* To ensure success of the dialysis, Lucifer yellow (LY) was added to the pipette-solution. Bar, 20µm. B) The insets show the typical current profiles of an astrocyte in the wildtype (top) and knockout group (bottom) clamped at -70 mV in response to 10 de- and hyperpolarizing voltage-steps in hippocampus. Only cells which displayed a series resistance of ≤ 125% of the initial value after 20 min of dialysis were included in the analysis. C) The graph shows the averaged current to voltage relationship of both genotypes at the start and the end of the 20 min dialysis period in hippocampus (black: wildtype, grey: BT-IgSF knockout). D and E) The boxplot graphs compare the membrane capacitance and membrane resistance of hippocampal astrocytes at the start and the end of the dialysis. No significant differences were observed in either comparison (2-way ANOVA, p > 0.05). Similar results on the current profiles, membrane capacitance and membrane resistance were obtained from cortical astrocytes.

**Extended Figure 6-1.**
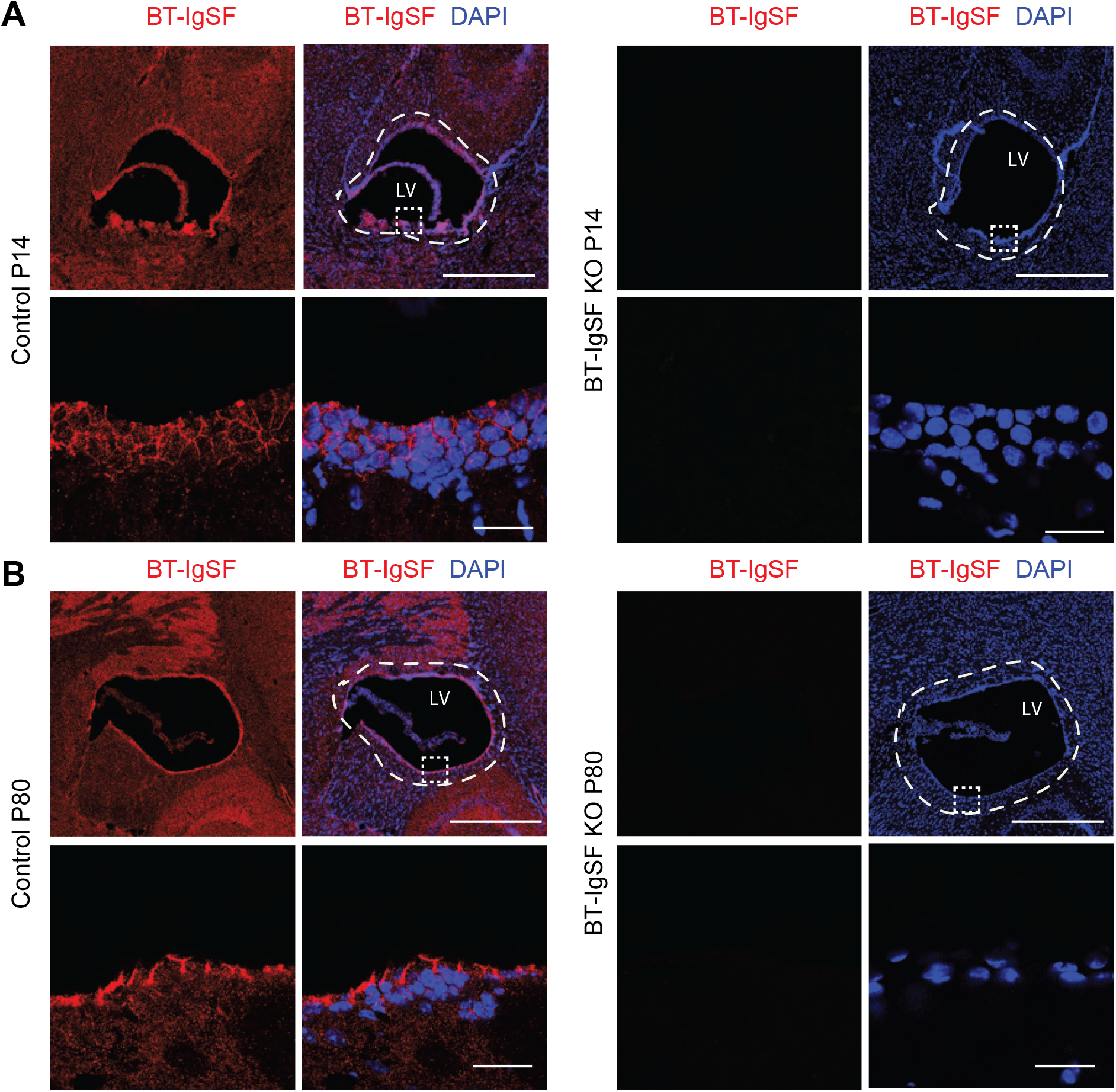
Localization of BT-IgSF at the lateral ventricle at P14 and P80. A and B) Coronal sections of the brain at P14 and P80 were stained with rabbit anti-BT-IgSF. BT-IgSF is strongly localized in cells lining the lateral ventricle. The dashed boxes indicate the position shown in row two or four. Absence of staining in BT-IgSF knockout tissues indicates the specificity of antibodies to BT-IgSF. LV, lateral ventricle. Scale bar, first and third row, 1 mm; scale bar second and fourth row, 20 µm.

**Table S1:** Antibodies for immunohistochemistry, immunocytochemistry and for Western blotting.

| Antibody | IHC and ICC | Western blots | Source |
| --- | --- | --- | --- |
| mAb γ-adaptin | 1 μg/ml | | BD Transduction Laboratories #610385 |
| Rabbit anti-BT-IgSF-Fc (Rb95 or Rb96) IgG | 1-3 μg/ml | 1 μg/ml | Pelz et al., 2017 |
| Rabbit anti-CAR | 1 μg/ml | | Patzke et al., 2010 |
| mAb anti-connexin43 | 1 μg/ml | | BD Transduction Laboratories, #610061 |
| Rabbit anti-connexin43 | 1:200 | 1:1000 | Cell signaling, #3512 |
| Rabbit anti-connexin30 | 1 μg/ml | 0.5 μg/ml | Invitrogen #71-2200 |
| Rabbit anti-connexin36 |  | 1 µg/ml | Zymed/Invitrogen 36-4600 |
| mAb connexin36 (H9) |  | 1 µg/ml | Santa Cruz sc-398063 |
| Rabbit anti-connexin36 (RbB5) | 0.5 µg/ml | 0.5 µg/ml | This study |
| mAb α-Tubulin |  | 0.025 µg/ml | Calbiochem CP06 |
| mAb anti-neurofilament | 7.5 µg/ml |  | mAb 2H3 Developmental Hybridoma Bank |
| mAb connexin35/36 |  | 1 µg/ml | Millipore MAB3045 |
| mAb ZO-1 | 1:200 |  | Invitrogen #339100 |
| Guinea pig anti-GFAP | 1:500 | 1:1000 | Synaptic Systems#173004 |
| Goat anti-GFAP | 1:200 | 1:1000 | Santa Cruz #sc-6170 |
| Rabbit anti-GFAP | 1:1000 |  | Dako Z0334 |
| mAb GM130 | 1 µg/ml |  | BD Transduction Laboratories #610823 |
| mAb LAMP-1 (1D4B) | 1 µg/ml |  | Santa Cruz ,sc-19992 |
| mAb anti-MAP2a/b | 1:500 | 1:1000 | Dianova#DLN-08583 |
| mAb anti-ZO-1 | 5 µg/ml |  | Invitrogen #339100 |
| Rabbit anti-ZO-1 | 1:100 |  | Invitrogen #40-2200 |
| mAb anti-GAPDH (1D4) |  | 0.5 µg/ml | Novus Biologicals, #NB300-221 |
| mAb anti-Clathrin (heavy chain) |  | 1:1000 | BD Transduction Laboratories #610499 |
| Goat anti-Rabbit-Cy3 | 1:400 – 1:1000 |  | Dianova |
| Goat anti-Mouse-Alexa488 | 1:400 – 1:1000 |  | Molecular Probes |
| Rabbit anti-Goat-Alexa488 | 1:500 |  |  |
| Goat anti-Rabbit-HRP |  | 1:20 000 | Dianova |
| Goat anti-Mouse-HRP |  | 1:20 000 | Dianova |

